# High-throughput discovery of trafficking-deficient variants in the cardiac potassium channel *KCNH2*: Deep mutational scan of *KCNH2* trafficking

**DOI:** 10.1101/2020.02.17.952606

**Authors:** Krystian A. Kozek, Andrew M. Glazer, Chai-Ann Ng, Daniel Blackwell, Christian L. Egly, Loren R. Vanags, Marcia Blair, Devyn Mitchell, Kenneth A. Matreyek, Douglas M. Fowler, Bjorn C. Knollmann, Jamie Vandenberg, Dan M. Roden, Brett M. Kroncke

**Author notes:** These authors contributed equally (Krystian A. Kozek, Andrew M. Glazer). Correspondence should be addressed to Brett M. Kroncke, PhD, Vanderbilt University Medical Center, 2215B Garland Ave, 1225E MRBIV, Nashville, TN 37232.

## Abstract

**Background:** *KCHN2* encodes the K_V_11.1 potassium channel responsible for *I_Kr_*, a major repolarization current during the cardiomyocyte action potential. Variants in *KCNH2* that decrease *I_Kr_* can cause Type 2 Long QT syndrome, usually due to mistrafficking to the cell surface. Accurately discriminating between variants with normal and abnormal trafficking would help clinicians identify and treat individuals at risk of a major cardiac event. The volume of reported non-synonymous *KCNH2* variants preclude the use of conventional electrophysiologic methods for functional study.

**Objective:** To report a high-throughput, multiplexed screening method for *KCNH2* genetic variants capable of measuring the cell surface abundance of hundreds of missense variants in *KCNH2*.

**Methods:** We develop a method to quantitate *KCNH2* variant trafficking on a pilot region of 11 residues in the S5 helix, and generate trafficking scores for 220/231 missense variants in this region.

**Results:** For 5/5 variants, high-throughput trafficking scores validated when tested in single variant flow cytometry and confocal microscopy experiments. We additionally compare our results with planar patch electrophysiology and find that loss-of-trafficking variants do not produce *I_Kr_*, but that some variants which traffic normally may still be functionally compromised.

**Conclusions:** Here, we describe a new method for detecting trafficking-deficient variants in *KCNH2* in a multiplexed assay. This new method accurately generates trafficking data for variants in *KCNH2* and can be readily extended to all residues in Kv11.1 and to other cell surface proteins.

**CLINICAL IMPLICATIONS:** Hundreds of *KCNH2* variants have been observed to date, and thousands more will be found as clinical and population sequencing efforts become increasingly widespread. The major mechanism of K_V_11.1 loss of function is misfolding and failure to traffic to the cell surface. Deep mutational scanning of *KCNH2* trafficking is a scalable, high-throughput method that can help identify new loss of function variants and decipher the large number of *KCNH2* variants being found in the population.

## INTRODUCTION

*KCNH2*, also known as the human Ether-à-go-go-Related Gene (hERG), encodes the rapid component of the delayed inward-rectifying, voltage-gated, cardiac potassium channel K_V_11.1. This channel produces a major repolarizing current in cardiomyocytes, *I_Kr_*. *KCNH2* loss-of-function variants are the second most common cause of congenital long QT syndrome (LQT2; MIM: #613688, Figure 1A).^1^ *KCNH2* gain-of-function variants are the most common cause of short QT syndrome (SQTS; MIM: #609620).^2^ As the names suggest, LQT2 is characterized by a prolonged QT interval on the electrocardiogram, and SQTS, a rarer disease, is characterized by a short QT interval. Both LQT2 and SQTS predispose individuals to syncope and sudden death.^2, 3^ To date, more than 400 *KCNH2* variants have been observed in patients with arrhythmias or sudden death, and over 500 *KCNH2* variants are present in the Genome Aggregation database (gnomAD), a large database of presumably mostly unaffected individuals.^1, 4–9^ With the onset of large population sequencing projects and increasing use of clinical genetic testing, the number of observed *KCNH2* variants is rapidly growing, much faster than the detailed characterization of these variants.^10^ An important challenge for *KCNH2* variant annotation is the identification and characterization of potentially disease-causing *KCNH2* variants found in individuals with or without a clinical phenotype.

**Figure 1:**
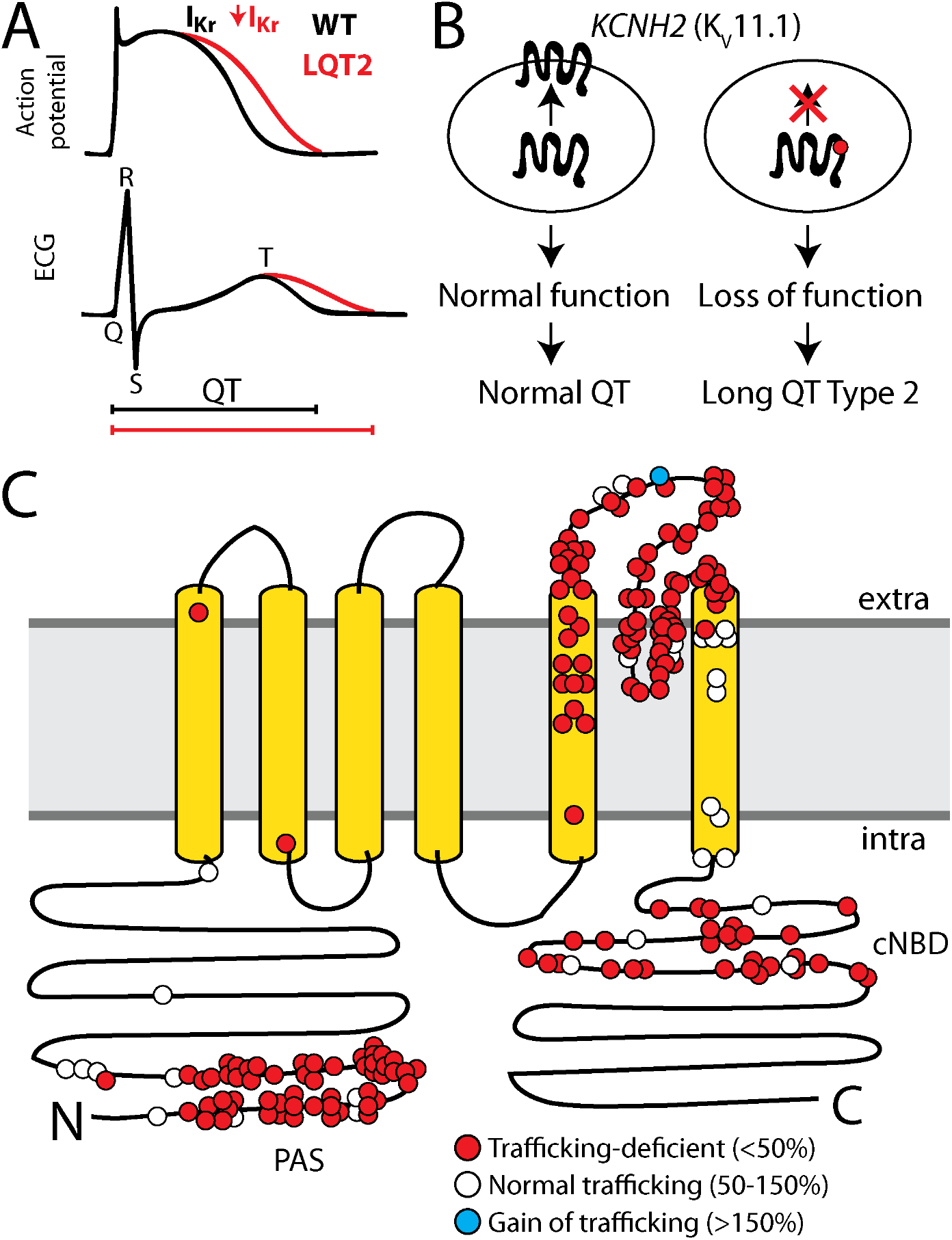
KCNH2 trafficking deficiency and Long QT Syndrome. A) *KCNH2* variants can cause Long QT Syndrome Type 2 (LQT2) by reducing *I_Kr_* inward potassium current. LQT2 manifests as a prolonged action potential in cardiomyocytes and a prolonged QT interval on the electrocardiogram. B) Schematic of K_V_11.1 trafficking to the plasma membrane. A common mechanism of LQT2 is reduced trafficking of K_V_11.1 to the cell surface. C) For missense variants, trafficking-deficiency is the dominant mechanism of LQT2 (178/210 variants; 85%). LQT2 variants that have been assessed for cell surface abundance are labeled in red (trafficking-deficient; <50% of wildtype) or white (trafficking-normal; 50-150% of wildtype). A Short QT Syndrome-associated variant, N588K, is labeled in blue; this variant has increased cell surface abundance (>150%). PAS indicates the approximate location of the PAS (Per-Arnt-Sim) domain thought to play a role in cell surface expression, cNBD indicates the Cyclic Nucleotide Binding Domain.

K_V_11.1 is an 1,159 residue protein with six membrane-spanning helices that forms a homotetrameric voltage-gated potassium channel at the plasma membrane surface in cardiomyocytes. A very common mechanism by which pathogenic variants cause protein dysfunction is destabilization-induced misfolding.^11–14^ For K_V_11.1, misfolding most commonly causes loss of trafficking to the plasma membrane, resulting in fewer functional channels at the cell surface (Figure 1B).^15–17^ 178/211 (84%) *KCNH2* variants associated with LQT2 whose cell surface abundance has been assayed have defective trafficking to the cell surface (Figure 1C).^18–28^

Here, we develop a deep mutational scanning (DMS) method that combines saturation mutagenesis with a high-throughput flow-cytometry-based trafficking assay. This approach offers a unique opportunity to generate high-throughput experimental data for hundreds or thousands of single codon substitutions in a multiplexed experiment. This provides complementary information to *in silico* predictions^29, 30^ and other functional characterization techniques, such as automated patch clamp electrophysiology^31^ and CRISPR-generated, or naturally-occurring, variants in induced pluripotent stem cell-derived cardiomyocytes.^32^

Here we present a DMS of a transmembrane helix of *KCNH2*. We generate 220 variants in an 11 amino acid region of *KCNH2* and assay the impact of these variants on cell surface abundance with a multiplexed flow cytometry assay. We determine cell surface abundance scores for these variants, and find that these scores correlate strongly with ion channel topology. This method predicts new deleterious variants in *KCNH2* likely to induce LQT2 and can be expanded to the rest of *KCNH2* and other membrane proteins.

## METHODS

### KCNH2 *mutagenesis*

We modified a previously published promoterless dsRed-Express-derivative plasmid^33^ and inserted wildtype *KCNH2* (hERG 1a; ENST00000262186) immediately adjacent to a recombinase AttB site. A “tile” system was created to divide up the size of fragments digested during molecular biology experiments by using naturally-occurring restriction enzyme sites and generating additional sites by synonymous mutations, which were introduced by a Quikchange Lightning Multi kit (Agilent). In this way, we divided wildtype *KCNH2* into 5 tiles of 500-800 base pairs (Figure S1). A previously published^34^ 9 amino acid HA tag with surrounding 4 amino acid linkers (NSEH<u>YPYDVPDYA</u>VTFE) was inserted between amino acids T443 and E444 (in the extracellular region between the S1 and S2 helices) by inverse PCR^35^ (Table S1). This tag has been demonstrated previously not to affect the electrophysiological properties of the channel.^34^ The promoterless “wildtype” plasmid (AttB-*KCNH2*-HA:IRES:mCherry) used for stable line generation contained an AttB recombination sequence (for genome integration), wildtype *KCNH2*-HA, an internal ribosome entry site (IRES), an mCherry reporter, and a SV40 termination sequence. A barcode integration site was located adjacent to the attB site, which allowed tagging of variants with random DNA barcodes (see below). We also generated a single tile plasmid (Tile 3-KCNH2) containing only Tile 3 (c.1474-1966), which was cloned into the dsRed-Express-derivative plasmid, surrounded by BglII and NdeI restriction sites.

Comprehensive codon mutagenesis was performed by inverse PCR of the Tile 3-*KCNH2* plasmid with 1 primer pair per codon. Each forward primer contained 5’NNN (the nucleotide, N, is a mix of A/C/G/T), encoding all 64 possible codons.^35^ 11 PCR products (corresponding to the 11 mutated codons, residues 545-555; Figure S2A) were pooled, PCR purified, and phosphorylated with T4 Polynucleotide Kinase (New England Biolabs, NEB), ligated with T4 DNA ligase (NEB), and incubated with DpnI (NEB). The products were then PCR Purified (Qiagen) and electroporated into MegaX DH10B Electrocomp Cells (ThermoFisher) with a Gene Pulser Electroporation System (Bio-Rad). Plating experiments using various dilutions of the library onto LB-Ampicillin plates were performed to determine transformation efficiency and library diversity. The mutant Tile 3 plasmid pool was then subcloned into the AttB-KCNH2-HA:IRES:mCherry plasmid by restriction digest with BglII and NdeI (NEB), ligated with T4 ligase (NEB), and electroporated as described above. An 18-mer poly-N barcode was created by annealing primers (Table S1) and extended to make fully double stranded with Klenow polymerase (NEB). The barcode was then PCR purified (Qiagen) and inserted adjacent to the AttB site by restriction digest with BsiWI and XbaI (NEB), and the library was electroporated as described above. At each cloning step, selected colonies were Sanger sequenced to verify the success of each step and to ensure sufficient variant library and barcode diversity.

### Subassembly to link barcodes and mutants

To link barcodes to mutants, the library was digested with NdeI (NEB), gel extracted (Qiagen), and ligated with T4 ligase (NEB), which caused intramolecular ligation to bring the barcode adjacent to the mutated Tile 3 (Table S1). PCR was performed with Q5 polymerase (NEB) following manufacturer’s instructions with 10 PCR cycles (primers listed in Table S1). Samples were purified with Ampure XP beads (Beckman Coulter) following manufacturer’s instructions, verified by Bioanalyzer and qPCR, and sequenced on an Illumina NovaSeq instrument with 150 base paired end sequencing. Reads were analyzed with a custom python script to identify barcodes where over 80% of reads associated with a single mutation.

### Generation of stable cell lines

HEK293 cells were grown in alpha MEM (#15-012-CV, Corning) with 10% FBS, 1% GlutaMax (Gibco), and in an incubator at 37 °C with 5% CO_2_. The library was integrated into a previously published HEK293T “landing pad” cell line.^33^ This cell line contains a landing pad safe harbor location engineered to contain an AttP site (Bxb1 recombination site) between a tetracycline-inducible promoter and a Blue Fluorescent Protein gene (HEK TetBxb1BFP). Cells were grown to 40-60% confluency before transfection. On day 0, this cell line was transfected with a plasmid expressing Bxb1 integrase (pCAG–NLS–HA–Bxb1; Addgene #51271, a gift from Pawel Pelczar^36^) by transient transfection with FuGENE 6 (Promega). On day 1, the cells were transfected with the library of *KCNH2* variants using FuGENE 6. On day 6, cells were incubated in 1 µg/ml doxycycline in HEK media to induce expression from the landing pad tetracycline-sensitive promoter. The resulting cell lines that integrated the plasmid had a single variant expressed from the landing pad, along with mCherry. Non-integrated cells expressed tagBFP and were mCherry-negative.

### High-throughput quantitation of variant surface trafficking using cell sorting

The HEK293T-integrated library was sorted with fluorescence activated cell sorting (FACS) to obtain cells that had successfully integrated an AttB-containing plasmid. 24 hours after doxycycline was added, cells were harvested with TrypLE Express (Thermo Fisher), resuspended in HEK media, spun at 500g for 2 minutes, and resuspended in divalent-free PBS with 1% bovine serum albumin (BSA). Cells were filtered and mCherry-positive, BFP-negative cells were sorted with a BD Aria III instrument. Filters used for sorting were 535 nm (excitation) and 610/20 nm (emission) for mCherry and 405 nm (excitation) and 450/50 nm (emission) for BFP. Cells were sorted into PureCoat plates (Corning) to enhance cell recovery and recovered for at least 12 hours before moving to non-PureCoat plates. 1x Penicillin-Streptomycin (Gibco) was used in cell medium for at least 48 hours after cell sorting to protect from infection, after which antibiotic-free medium was again utilized.

After sorted cells were recovered and grown to confluency, a surface staining and fluorescence-activated cell sorting assay was used to assay cell surface abundance of the library. Cells were harvested with TrypLE Express (Thermo Fisher), resuspended in HEK media, spun at 500g for 2 minutes, and resuspended in divalent-free PBS with 1% BSA. Cells were filtered and incubated in a 1:500 solution of mouse anti-HA antibody covalently bound to Alexa 647 (Cell Signaling 6E2, product #3444). Cells were incubated with the antibody by shaking vigorously for 15-30 minutes, then spun at 500g for 2 minutes. Pelleted cells were then resuspended in PBS with 1% BSA and filtered. A second cell sort was then performed. Cells that were both mCherry positive and tagBFP negative, indicating successful plasmid integration, were identified as described above. These cells were sorted into 4 groups depending on level of Alexa 647 signal (Figure S2). The Alexa 647(-) pool was defined based on the staining levels of control cells not treated with doxycycline. The remaining 3 pools were chosen to have approximately the same number of cells in each pool. The four pools of sorted cells were replated and grown to confluency as described above. After multiple days of growth to achieve >2 million cells per pool, cells were harvested using TrypLE Express (Thermo Fisher) and pelleted at 1 million cells/tube for frozen storage.

### Illumina sequencing of libraries

DNA was isolated from each of the four cell pools with 100 µL QuickExtract (Lucigen) per 1 million cells following the manufacturer’s instructions. PCR to amplify the barcode for Illumina sequencing was performed on 2.5 µL of isolated DNA (primers listed in Table S1) with Q5 polymerase (NEB), following manufacturer’s instructions with between 25-35 cycles. Illumina libraries were cleaned with AmpureXP beads (Beckman Coulter) following manufacturer’s instructions, assessed with Bioanalyzer and qPCR, and sequenced on an Illumina NovaSeq instrument with 150 base paired end sequencing. Sequencing reads were de-multiplexed and processed with custom python scripts to identify the barcode present in each read.

### Calculation of trafficking scores

For each barcode, a raw trafficking score was calculated as the weighted average of the abundance of each barcode (i) in the four pools:

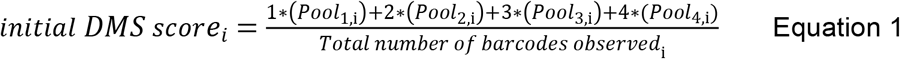

Where Pool_n,i_ is the fraction of the i^th^ barcode in the n^th^ pool of sorted cells; n ranges from 1 (no K_V_11.1 present, i.e. no AF647 signal) to 4 (high abundance of K_V_11.1, i.e. high AF647 signal). The scores were then aggregated by variant and normalized with a linear transformation so that trafficking score ranged from 0 (barcodes only observed in the AF647 negative pool) to 100 for WT. The scores were then averaged across the 2 replicate experiments (separate transfections of the mutant library pool).

### Flow cytometry, confocal microscopy, and patch clamping assay of individual *KCNH2* variants

Single variants were introduced by mutagenesis of the Tile 3-*KCNH2* plasmid (Table S1) using the Quikchange Lightning Multi kit (Agilent), with 1 primer per variant, following the manufacturer’s instructions. After the mutation was verified with Sanger sequencing, the mutant Tile 3 was subcloned into the full length AttB-KCNH2-HA:IRES:mCherry by restriction digest with BglII and NdeI (NEB) and ligation with T4 ligase (NEB). Plasmids were then verified with Sanger sequencing. Stable HEK293T landing pad cell lines were generated for each mutant as described above. For flow cytometry, cells were stained with an anti-HA antibody and assayed for cell surface abundance as described above. For confocal microscopy, live cells were first stained with an anti-HA antibody directly conjugated to Alexa 647 as described above. Then, cells were fixed for 10 minutes in 4% paraformaldehyde, washed in PBS, permeabilized in 0.2% Triton-X, washed in PBS, and incubated in an anti-HA antibody directly conjugated to Alexa 488 (Cell Signaling #2350) for 1 hour. Next, cells were washed thrice in PBS and incubated in 1:1000 DAPI. Imaging was performed using an LSM 880 inverted laser scanning confocal microscope with a 63X/1.40 PlanAPOCHROMAT oil immersion objective. Each channel was scanned sequentially to minimize bleed through and the emission was collected using photomultiplier tube detectors. DAPI was excited using a 405 nm diode laser and the emitted light was collected from 410-490 nm. Alexa Fluor 488 was excited using a 488 nm argon laser and collected from 500-544 nm. mCherry was excited using a diode-pumped solid state 561 nm laser and collected from 562-624 nm. Alexa Fluor 647 was excited at 633 nm and collected from 638-755 nm. The pinhole was set to produce an optical slice depth of 1.3 µm.

#### Automated patch clamp recording

Automated patch clamp recordings were performed in the whole cell voltage clamp configuration using medium resistance (4–4.5 MΩ) single hole chips at ∼25 °C using a SyncroPatch 384PE platform (Nanion Technologies, Munich, Germany). Data were acquired at a sampling rate of 5 to 10 kHz, depending on the protocol, using PatchControl384 v1.5.6 software (Nanion Technologies). External recording solution contained (in mM): NaCl 140, KCl 5, CaCl_2_ 2, MgCl_2_ 1, HEPES 10, D-Glucose 5; adjusted to pH 7.4 with NaOH. Cell sealing was aided using a modified external solution containing 10 mM CaCl_2_. The internal solution contained (in mM): KF 110, KCl 10, NaCl 10, HEPES 10, EGTA 10; adjusted to pH 7.2 with KOH. Cells were detached by incubating with Accumax (Sigma-Aldrich, USA) for 5 minutes at 37 °C followed by centrifugation at 200 g for 3 minutes before resuspended in divalent free solution (in mM: NaCl 140, KCl 5, HEPES 10, D-Glucose 5; adjusted to pH 7.4 with NaOH). Cells were then incubated for 60 min at 10 °C while shaking on a rotating platform at 500 rpm, before being dispensed into the 384 wells of the patch engine.

#### Data analysis of patch clamp recording

To measure the voltage dependence of activation, cells were depolarized to a range of potentials (from –50 mV to +70 mV in 10 mV increments) for 1 s before stepping to –120 mV to record tail currents. The mid-point of the voltage dependence of activation (V_0.5_) was determined by fitting a Boltzmann distribution to a plot of the peak tail currents against the voltage of the depolarizing step, using Equation 2:

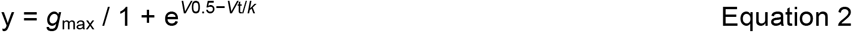

where: *V*_0.5_ is the half-maximum activation voltage, *V*t is the test potential, *k* is the slope factor and *g*_max_ is the maximum conductance.

To measure the rate of hERG channel closure (deactivation), cells were depolarized to +40 mV for 1 s before stepping to a range of tail potentials (from +20 to −150 mV in 10 mV increments) for 3 s. The decay component, after the initial ‘hooked’ current, represents K_V_11.1 channel deactivation (transition from open to closed state). To determine the time constant of deactivation, tail currents were fit with a double exponential function using Clampfit v11 software (Molecular Devices, San Jose, CA, USA). The overall time constant of deactivation was calculated as a weighted sum of the two components (i.e. fast and slow components of deactivation) recorded at –50 mV, as previously described,^31^ see Equation 3:

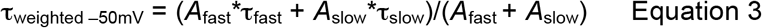

where *A* is the current amplitude and τ is the time constant. Data were analyzed and plotted using a combination of software, including the Microsoft Excel 365 (Microsoft, Redmond, WA, USA), DataController384 V1.5.2 (Nanion Technologies, Munich, Germany) and Prism 8 (GraphPad Software, San Diego, CA, USA). Recordings that satisfied the following quality control criteria were used: Seal resistance ≥ 0.3 GΩ; series resistance ≤20 MΩ; cell capacitance between 5 to 50 pF and the leak current is less than 40pA measured at –120mV. Whole-cell currents were normalized for membrane capacitance and presented as violin plots with median values as solid lines and upper and lower quartiles shown as dashed lines. Statistical significance was tested using Mann-Whitney tests.

## RESULTS

We modified a flow cytometry-based cell-surface trafficking assay^37^ for compatibility with a DMS of *KCNH2*. To accomplish this, we inserted an hemagglutinin (HA) tag^34^ into the first extracellular loop of *KCNH2*, overexpressed this plasmid in HEK293 cells, and isolated cells using flow cytometry.^33^ Control variants of *KCNH2* were generated within the HA-*KCNH2* construct, and trafficking characteristics were validated with flow cytometry and confocal microscopy (Figure 2A). Wildtype HA-*KCNH2* had a high surface abundance as detected by both flow cytometry or confocal microscopy (Figure 2B). In contrast, G601S and A561V, which have previously been described to have mistrafficking-related loss of function defects,^38, 39^ had reduced cell surface abundance with both methods (Figure 2B).

**Figure 2:**
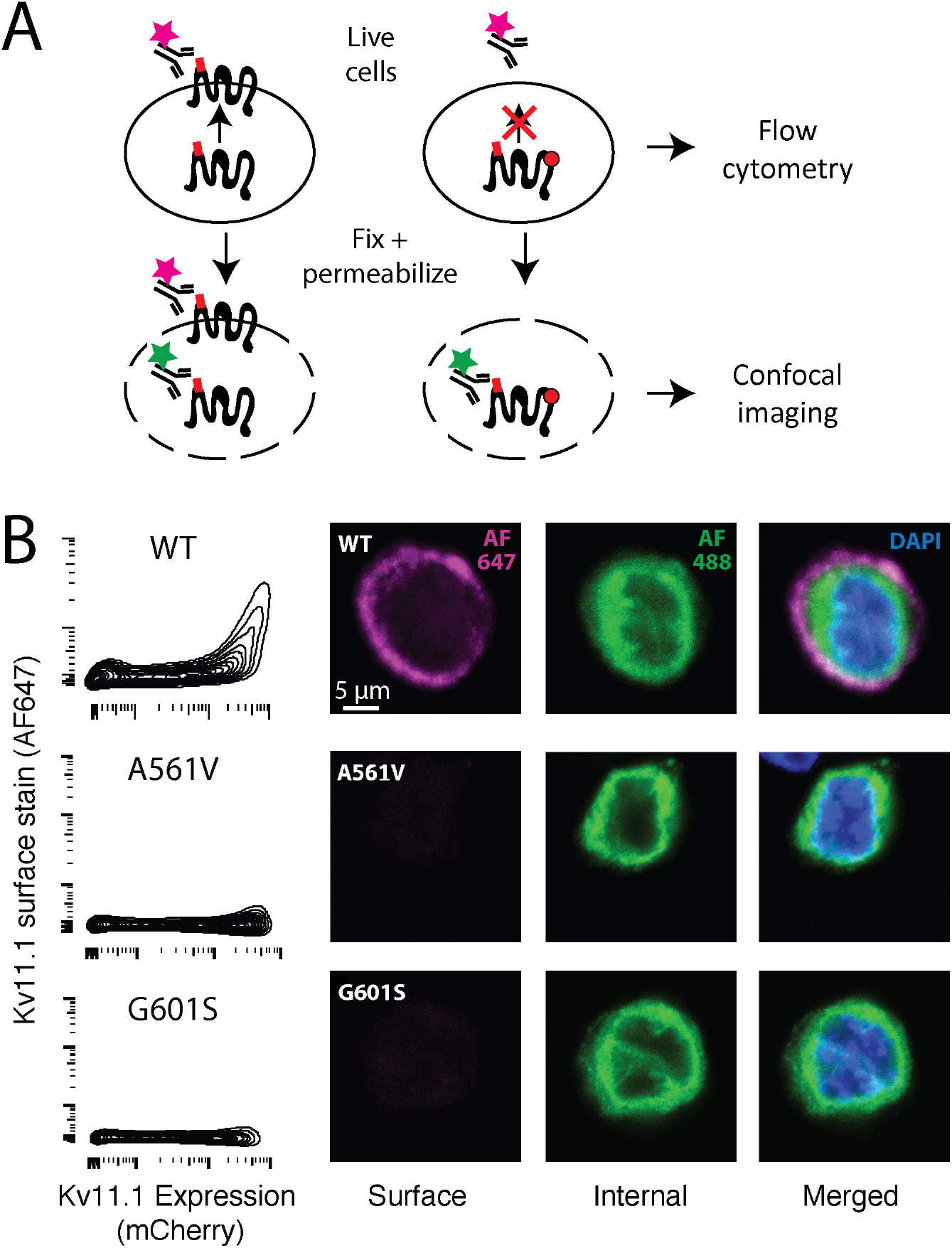
KCNH2 cell-surface trafficking assay. A) Diagram of antibody labeling of surface or surface and internal K_V_11.1. Live cells are stained with Alexa 647-labeled anti-HA antibody, which labels surface K_V_11.1. Cells are assayed by flow cytometry. For confocal imaging, cells are fixed, permeabilized, and stained with an Alexa 488-labeled anti-HA antibody, which labels internal K_V_11.1. B) Flow cytometry assay of HEK293 cells expressing wildtype *KCNH2*, G601S (partial loss of function), or A561V (near total loss of function). The x-axis indicates mCherry level, which is a marker of protein expression. C) Confocal imaging of HEK293 cells expressing wildtype *KCNH2*, G601S, or A561V.

### Deep mutational scan of 11 amino acid region of KCNH2

We next generated a barcoded plasmid library of nearly every possible variant in an 11 amino acid region (residues 545-555) of *KCNH2* using PCR with degenerate primers^35^ (Table S1). After subassembly^40^ to link barcodes to variants, 14,466 unique barcodes were detected, which included 220/231 possible missense variants (Figure S3). The 11-amino acid experiment successfully distinguished between two control classes of variants, nonsense variants (predicted to have no channel function) and synonymous variants (predicted to have wildtype channel function; Figure 3A). Initial trafficking scores were calculated from a weighted average of each variant’s abundance in the four sorted pools. These scores were averaged across two independently transfected and sorted replicate experiments. From a range of possible initial trafficking scores between 1 and 4, the nonsense variants had an average raw score of 1.09, whereas WT had an average score of 2.7 (Equation 1). We generated normalized scores such that 0 was the lowest possible score and 100 was the average score for WT (Figure 3). The missense scores ranged widely from loss of function to wildtype-like, and the mean score for all missense variants was 54.8 (Figure 3A and B). Trafficking scores were highly consistent across the two replicate experiments (Spearman’s r = 0.77).

**Figure 3:**
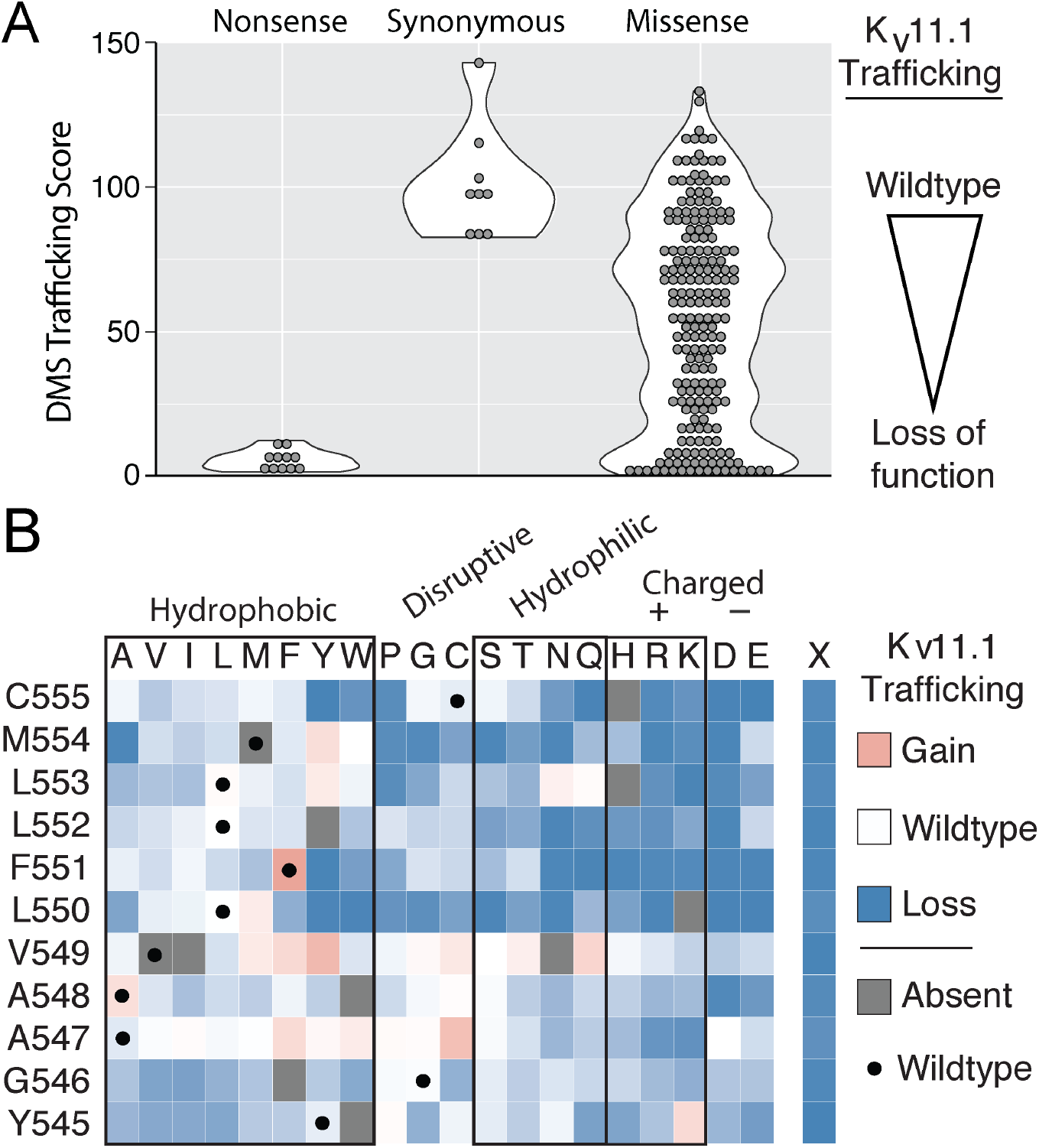
Deep mutational scan identifies trafficking-defective variants. A) The 11-residue pilot saturation mutagenesis experiment. Trafficking scores between nonsense and synonymous variants were highly separated, and were used to determine “wildtype” and “loss of function” levels and calculate normalized scores, which are shown in the plot. The missense variants had a range of scores from near-wildtype to loss-of-function. B) The wildtype residue is indicated in the leftmost column and amino-acid substitution is indicated on the top row. Blue indicates that the variant did not express to the plasma membrane, white indicates the variant traffics like WT, and red indicates the variant traffics more than WT. Out of the 220 variants assayed, 86 were WT-like (score > 75), 57 were mild loss-of-trafficking (75 > score > 50), 48 were loss-of-trafficking (50 > score > 25), and 73 were severe loss-of-trafficking (score < 25). Note the low trafficking scores of charged or polar amino-acids closer to the middle of the membrane (residues 550-555) (see Figure 4).

### Trafficking scores and K_V_11.1 topology

We explored the structural basis of the trafficking scores with a three-dimensional structure of K_V_11.1 (Figure 4).^41^ Residues Y545-V549 tolerated substitutions more than residues L550-C555, especially secondary structure-disruptive substitutions (glycine and proline) and hydrophilic substitutions (Figure 3B). These observations are consistent with the topology and position within the membrane of the pilot region: residues 545-549 join the S4 and S5 helices on the intracellular membrane leaflet, a segment transitioning from random coil to α-helical secondary structure (Figure 4). Missense substitutions at residues 545-549 were less likely on average to induce mistrafficking compared to residues 550-555, a mean trafficking score of 70.2 versus 42.2, respectively. In contrast, residues 550-555, closer to the membrane interior, were less tolerant to any substitution, especially polar and charged residues. The trafficking scores correlated modestly with computational predictions, with a Spearman’s ρ of 0.31 and 0.21 with PROVEAN and PolyPhen-2, respectively (File S1).

**Figure 4.**
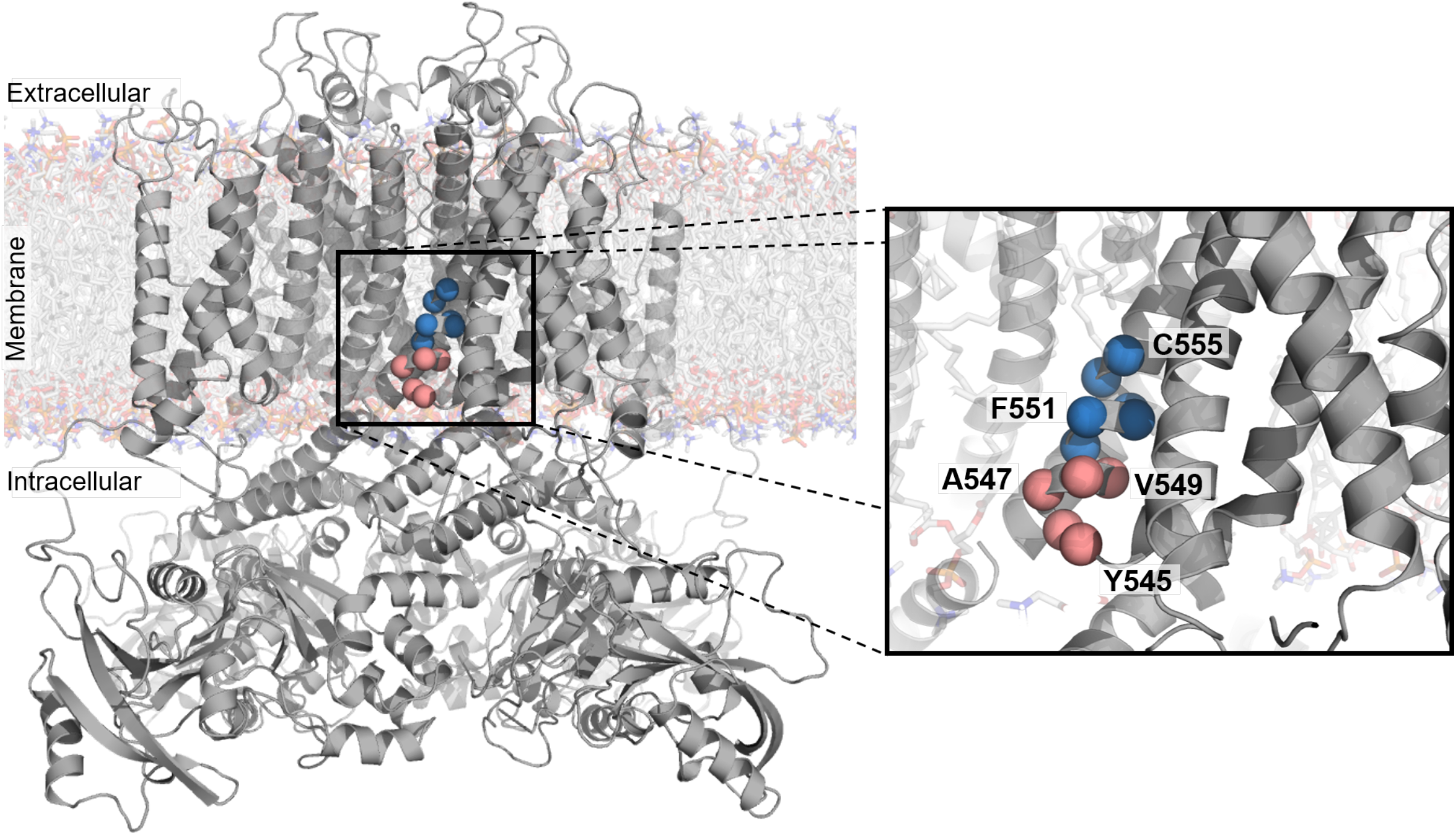
Polar and charged residues close to the middle of the membrane cause trafficking-deficiency. *In silico* model of the protein product of KCNH2 (K_V_11.1) with residues mutagenized in a pilot study of a high-throughput trafficking assay highlighted as red or blue spheres. These residues are on the intracellular half of the S5 transmembrane helix. The red to blue transition from V549 to L550 reflects the transition from residues tolerant of hydrophilic substitution to residues intolerant of hydrophilic substitution, respectively, as observed in the high-throughput trafficking assay (Figure 3B).

### Validation of trafficking assay scores

To validate our trafficking scores, we selected five variants for further characterization. These variants were selected based on their unintuitive DMS trafficking results: L552N and A547D are in regions intolerant of hydrophilic substitution yet had a trafficking score like WT (scores of 108 and 99, respectively); M554A is in a region tolerant of hydrophobic substitution yet does not traffic (score of 4); and M554W is a disruptive substitution yet traffics like WT (score of 99), in contrast to M554A. F551Y. In single variant validation experiments, these five variants had confocal and flow cytometry cell surface abundance concordant with their DMS scores (Figure 5 and Table 1). All variants in the validation set are predicted to be deleterious by PROVEAN and PolyPhen-2.

**Figure 5:**
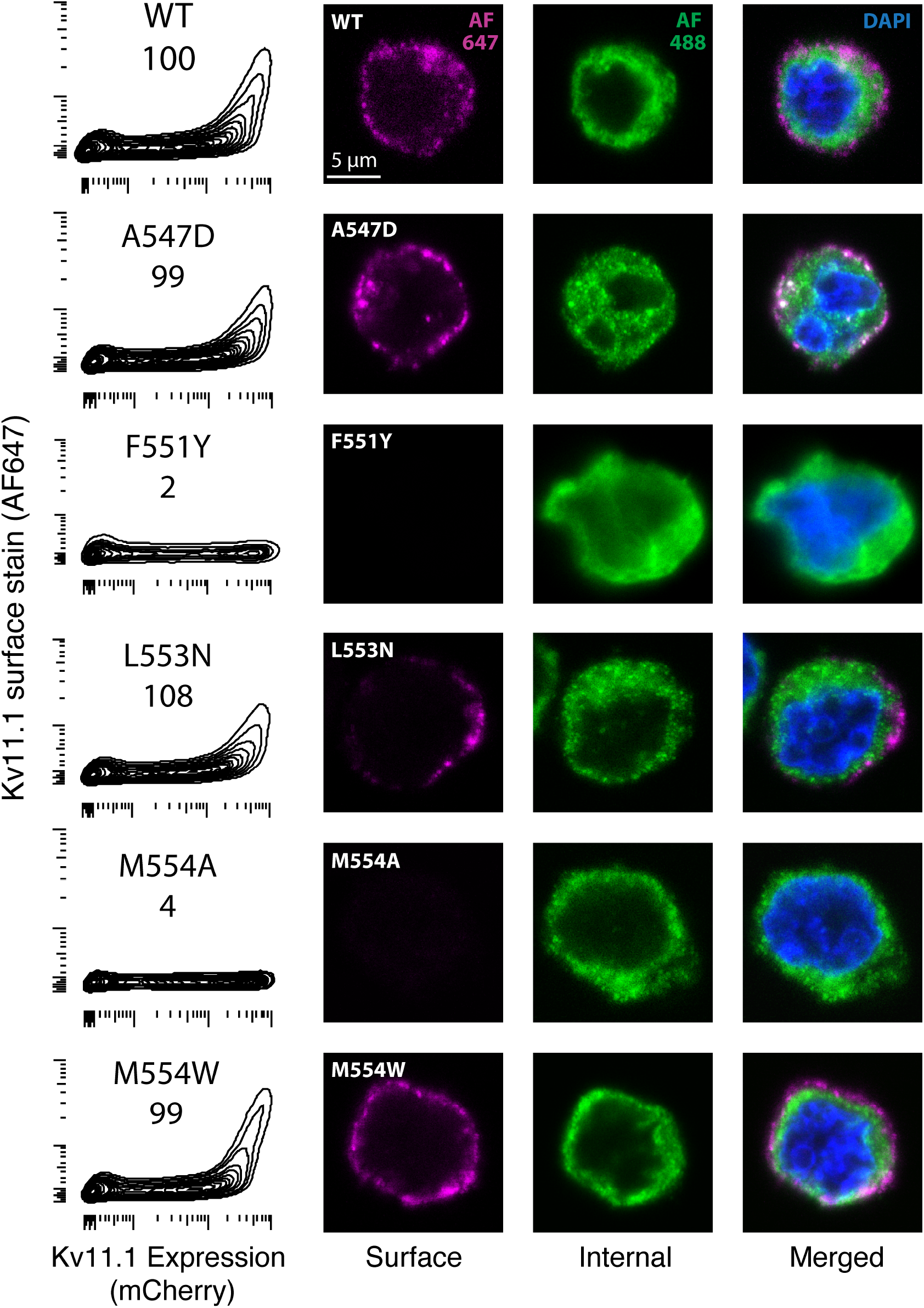
Validation of trafficking scores. Validation of selected variants by analytical flow cytometry (left) and confocal microscopy (right). In the leftmost column, DMS score is indicated directly below variant name. X-axis is mCherry fluorescence (indicating K_V_11.1 expression), the y-axis is surface stained K_V_11.1 (protein product of *KCNH2*). The remaining columns are extracellularly stained K_V_11.1 (AF647, purple) intracellularly stained K_V_11.1 (AF488, green), and both merged with DAPI stained nucleus (blue).

**Table 1:** Trafficking score validation.

| Variant | DMS score | Flow/Confocal validation | Patch clamp |
| --- | --- | --- | --- |
| A547D | 99 | WT-like | LOF |
| L553N | 108 | WT-like | Fast deactivation |
| M554A | 4 | No trafficking | LOF |
| M554W | 99 | WT-like | LOF |
| F551Y | 2 | No trafficking | -- |

PROVEAN and PolyPhen-2 predict all variants listed are deleterious (File S1)

In addition, we examined four variants in the studied region that have previously been observed in literature reports or individuals in the gnomAD database.^9^ L552S was previously reported as a loss of trafficking variant^15^ and has been observed in 90 cases of Long QT syndrome in the literature, 28 unaffected individuals, and 20 alleles in the gnomAD database. Our DMS score for L552S was 25, consistent with the strong association with LQT2 and a loss of trafficking as previously reported.^15^ Three other variants, A547T, V549M, and F551L, were not observed in any literature cases of Long QT Syndrome and have been observed in 1-2 cases in the gnomAD database, too few to assess relevance from patient data alone. All of these variants had normal trafficking scores (Table S2). Thus, the DMS trafficking scores are consistent with the available data for variants in this region.

To further assess the trafficking results, we collected planar patch clamp data on the five validation variants. Consistent with its loss of trafficking DMS score (score of 4) and its loss of trafficking in validation experiments, M554A also had a total loss of I_Kr_. Surprisingly, the other three validation variants, which had detectable trafficking by DMS and in validation experiments, also had loss of function defects by patch clamp (Figure 6). Despite being present at the cell surface (Figure 5), M554W and A547D did not produce measurable *I_Kr_*, while L553N had detectable *I_Kr_*, but with lowered peak tail current and faster deactivation rate, essentially producing a loss-of-function phenotype (Figure 6). Therefore, the high-throughput trafficking scores were successful at identifying loss of function variants that acted through a trafficking mechanism, but not by other mechanisms.

**Figure 6:**
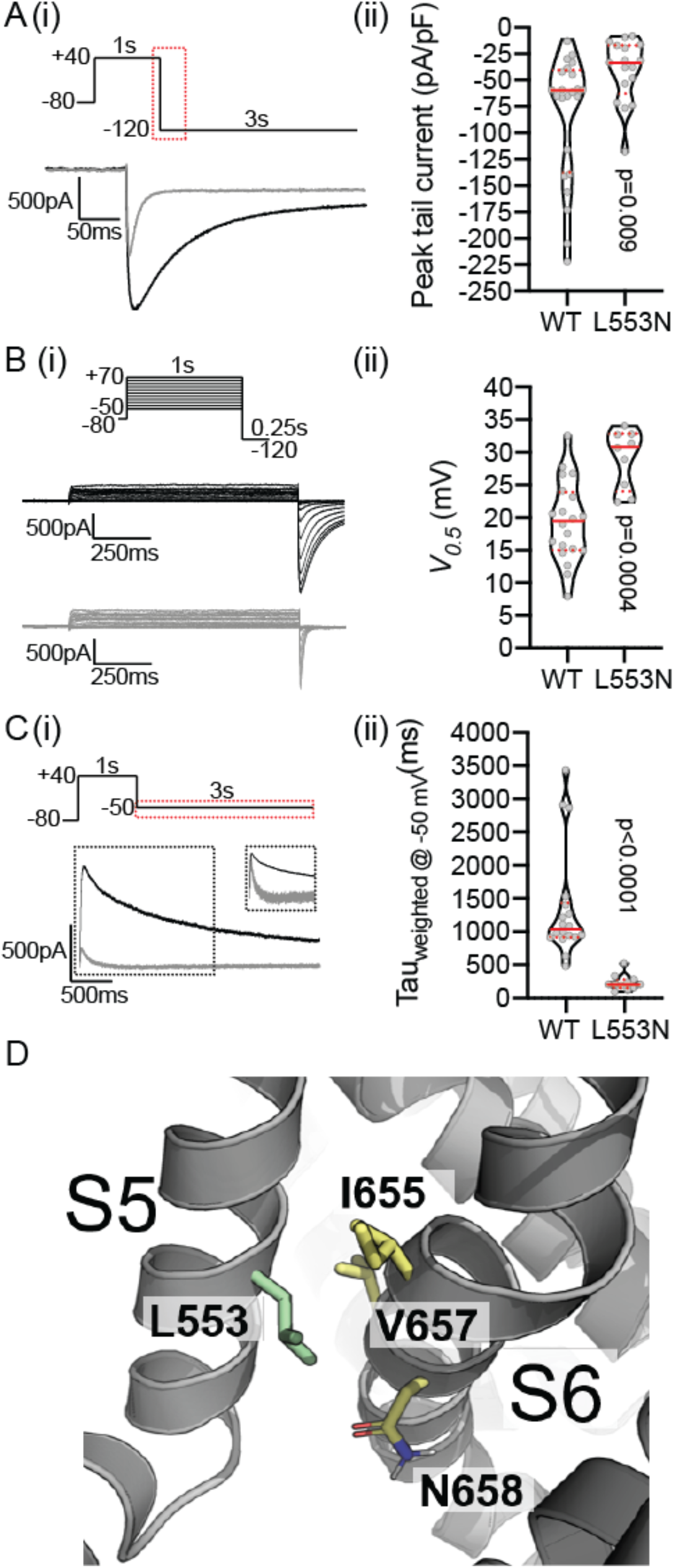
Functional characterization of L553N. Ai) All channels were fully activated at +40mV for 1s before stepping to –120mV for 3s to allow for recovery from inactivation and measure current density. The peak tail current traces for WT (black) and L553N (grey) are shown for the region highlighted in the voltage protocol. Aii) Violin plot of peak tail current amplitudes normalized to cell capacitance. Bi) Typical families of current traces recorded during Isochronal activation protocols, depolarizing the channels from – 50mV to +70mV for 1s to activate the channels before stepping to –120mV to record the tail currents, for WT (black) and L553N (grey). Bii) Violin plot of *V*_0.5_ values derived by fitting Boltzmann function to the tail current amplitudes (see Methods for details). Ci) The current traces for WT (black) and L553N (grey) were shown for the region highlighted in the voltage protocol after channels were fully activated at +40mV for 1s before stepping to –50mV for 3s to initiate deactivation. Cii) Violin plot of the weighted time constant (Τ_weighted_) for channel deactivation at –50mV, (see methods for detail of calculation of Τ_weighted_). In all violin plots, median and quartiles are shown as solid and dotted lines, respectively. The reported two-tailed P-values were derived from Mann-Whitney tests. D) Structural context of L553 (on helix S5), which points between residues V657 and N658 on the S6 helix within the same subunit.

## DISCUSSION

### A high-throughput assay for identifying *KCNH2* variant trafficking

We developed a scalable, multiplexed flow cytometry and high-throughput sequencing assay to measure the cell surface abundance of hundreds (and potentially thousands) of variants in *KCNH2*. We generated trafficking scores for 220/231 possible missense variants in an 11 amino acid region of *KCNH2*. We validated these scores in several ways. First, the scores correlated very well (Spearman ρ = 0.77) across the two replicate experiments. Second, the scores distinguished nonsense from synonymous variants (Figure 3). Finally, 5/5 missense variants with a range of surprising, unintuitive trafficking scores were validated by flow cytometry and confocal immunocytochemistry on single variants (Figure 5). Therefore, we believe these scores accurately capture cell surface abundance, which is the major mechanism for ∼80% of all LQT variants studied to date.^15–17^

### Improved understanding of *KCNH2* trafficking deficiency

From our trafficking data, we could observe two sub-domains in the 11-residue pilot region: a more exposed, hydrophilic region (residues 545-549) and a more buried hydrophobic region (residues 550-555). We find a difference in average trafficking behavior of variants in these two regions. In the hydrophilic region, hydrophilic substitutions were tolerated, whereas in the hydrophobic region, hydrophilic substitutions were more likely to lead to compromised trafficking. The tolerance of hydrophobic and hydrophilic substitutions in this pilot region additionally reflects the known topology of the *KCNH2* S5 helix, hydrophilic residues being less well-tolerated near the middle of the membrane bilayer (Figures 3B and 4). Our trafficking data have a Spearman’s ρ of 0.31 and 0.21 with PROVEAN and PolyPhen-2, respectively, suggesting modest overlap in information contained in each feature. L553N trafficked like WT and produced measurable peak tail current; however, this variant had a dramatically faster deactivation time, the time it takes the channel to stop conducting after the cell is repolarized (Figure 6). L553 points in between residues N658 and V659 in the S6 helix within the same subunit.^42^ Both residues are known to accelerate deactivation and the asparagine (N) substitution could stabilize an interaction between N658 and L553N resulting in a preferred deactivated state (Figure 6D).

### Approaches to studying *KCNH2* variants

This work represents a new tool to studying *KCNH2* variants in high-throughput. Several studies have previously quantified trafficking of K_V_11.1, typically using assays that rely on Western blotting or confocal imaging, which are not scalable to thousands of variants.^17, 43^ Recent advances in automated patch clamp electrophysiology have enabled the rapid characterization of dozens of voltage-gated potassium channel variants,^31, 44^ and induced pluripotent stem cell derived cardiomyocytes show promise to characterizing *KCNH2* variants in a non-heterologous system. However, the throughput of these methods does not allow for the rapid screening of the thousands of possible variants in *KCNH2*. In contrast, computational predictions like Polyphen2 or PROVEAN can be easily generated for every variant in *KCNH2*. However, while these computational predictions can partially predict disease risk, they are imperfect and often cannot capture the subtle differences among different nearby residues (File S1).

### Improving the prediction of *KCNH2* variant pathogenicity

Our assay has the potential to sample nearly all possible amino acid substitutions in *KCNH2* in an unbiased manner. This will provide a database for patients and clinicians to identify the effect a previously uncharacterized mutation may have on patient disease propensity. Future work will expand this method to much larger regions of *KCNH2*, with a focus on regions enriched for known LQT variants (Figure 1). Accomplishing this will allow a more comprehensive exploration of the relationship between *KCNH2* trafficking deficiency and disease risk. The method presented here is in principle generalizable to other cardiac ion channels, and all other proteins found at the plasma membrane, approximately 30% of the human proteome.^11–14^ Though protein destabilization-induced mistrafficking is the dominant etiology of variant pathogenesis across the proteome, our electrophysiological results highlight the need to approach trafficking data with caution. M554A does not traffic to the plasma membrane and it indeed yielded no current in patch clamp experiments, indicating the success of our screening approach. However, A547D and M554W trafficked normally in the high-throughput assay and in single variant validation experiments, yet did not produce *I_Kr_* currents in patch clamp experiments. This result indicates that certain variant loss of function is not through a trafficking mechanism, hence all variants with normal trafficking scores need to be interpreted with caution, as they may still cause loss of function through other mechanisms. Trafficking assays first will enable lower-throughput methodologies to focus on variants that are more amenable to surface-targeted functional studies. Future work may be possible that characterize *KCNH2* variant function in deep mutational scanning assays in addition to *KCNH2* surface abundance.^45^

#### Limitations

Seven variants were unable to be analyzed due to lack of barcoding during the generation of the *KCNH2* plasmid library. Additionally, our work was done in HEK cells and not cardiomyocytes. Therefore, we do not account for the expression of additional *KCNH2* splice variants and other subunits that may contribute to the inward rectifier current that underlies *I*_Kr_ in cardiomyocytes. Our assay investigates trafficking of homozygous *KCNH2* 1a variants with the most common allele (K897). Subsequent studies will investigate the effects of coexpression with WT splice isoforms, 1a and 1b, axillary subunits, e.g. *KCNE2*, and in the presence of common polymorphisms (K897T and R1047L).

#### Conclusion

We have developed a high-throughput trafficking assay to characterize cell surface expression of *KCNH2* variants in a massively parallel fashion. This method can be expanded to other ion channels and transmembrane proteins where altered cell surface abundance is a major mechanism of disease pathogenesis.

## Supporting information

File S1 DMS scores

## AUTHOR CONTRIBUTIONS

BMK conceived and designed the overall study. KAK conducted the high-throughput screen with the help of LRV and MB. AMG, BMK, KAM, and DMF generated reagents for performing the high-throughput screen. CN, DB, DM, LRV, and CLE performed validation experiments to characterize single variants. DMF, BCK, JV, and DMR provided guidance on experimental plans and data interpretation. All authors commented on the manuscript. KAK, AMG, and BMK co-wrote the manuscript.

## ACKNOWLEDGMENTS

We thank the following Vanderbilt core facilities: Vanderbilt Flow Cytometry Shared Resource, the Vanderbilt Cell Imaging Shared Resource, and the Vanderbilt VANTAGE genomics core.

## SUPPLEMENTAL MATERIAL

**Table S1.**
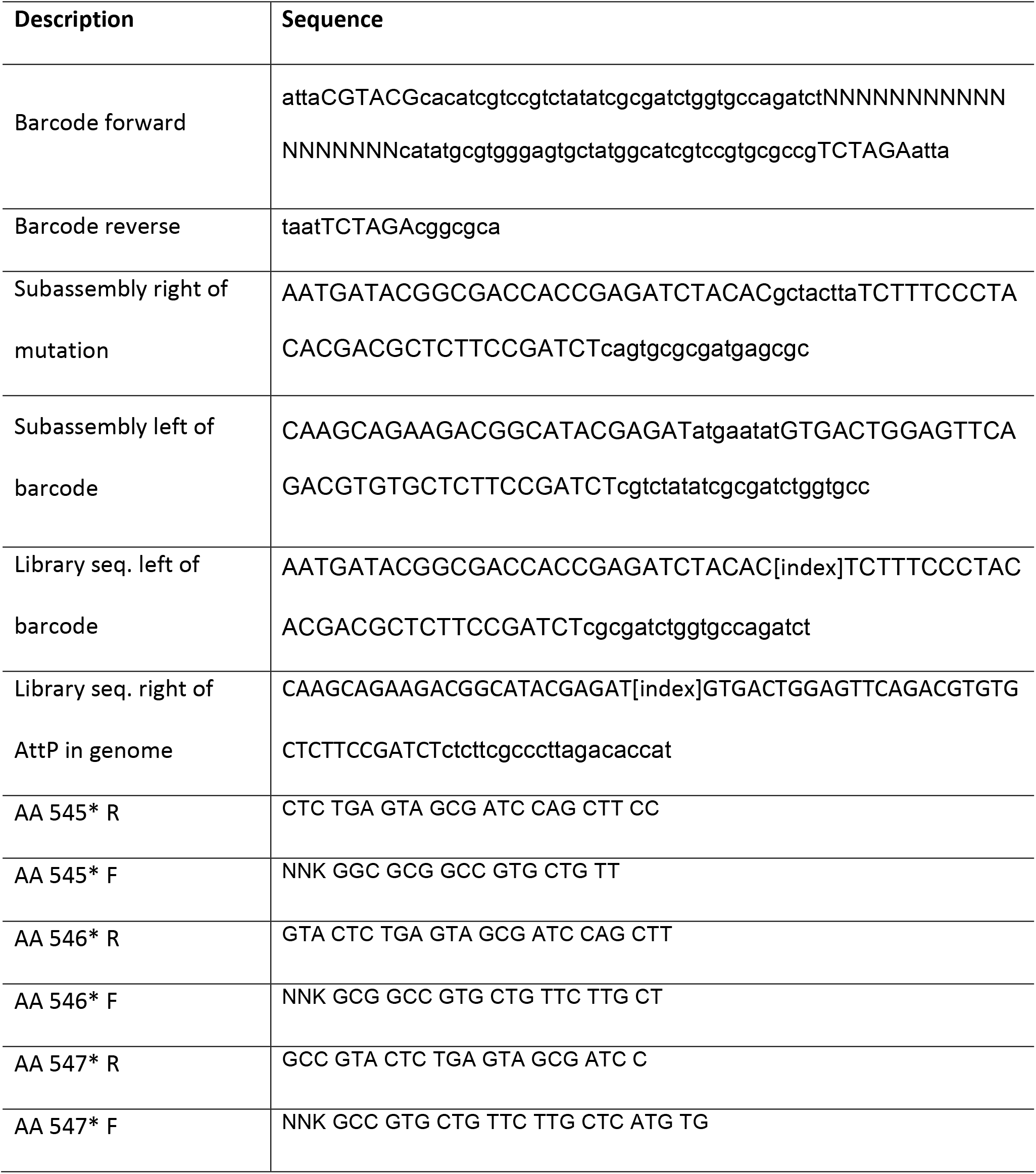
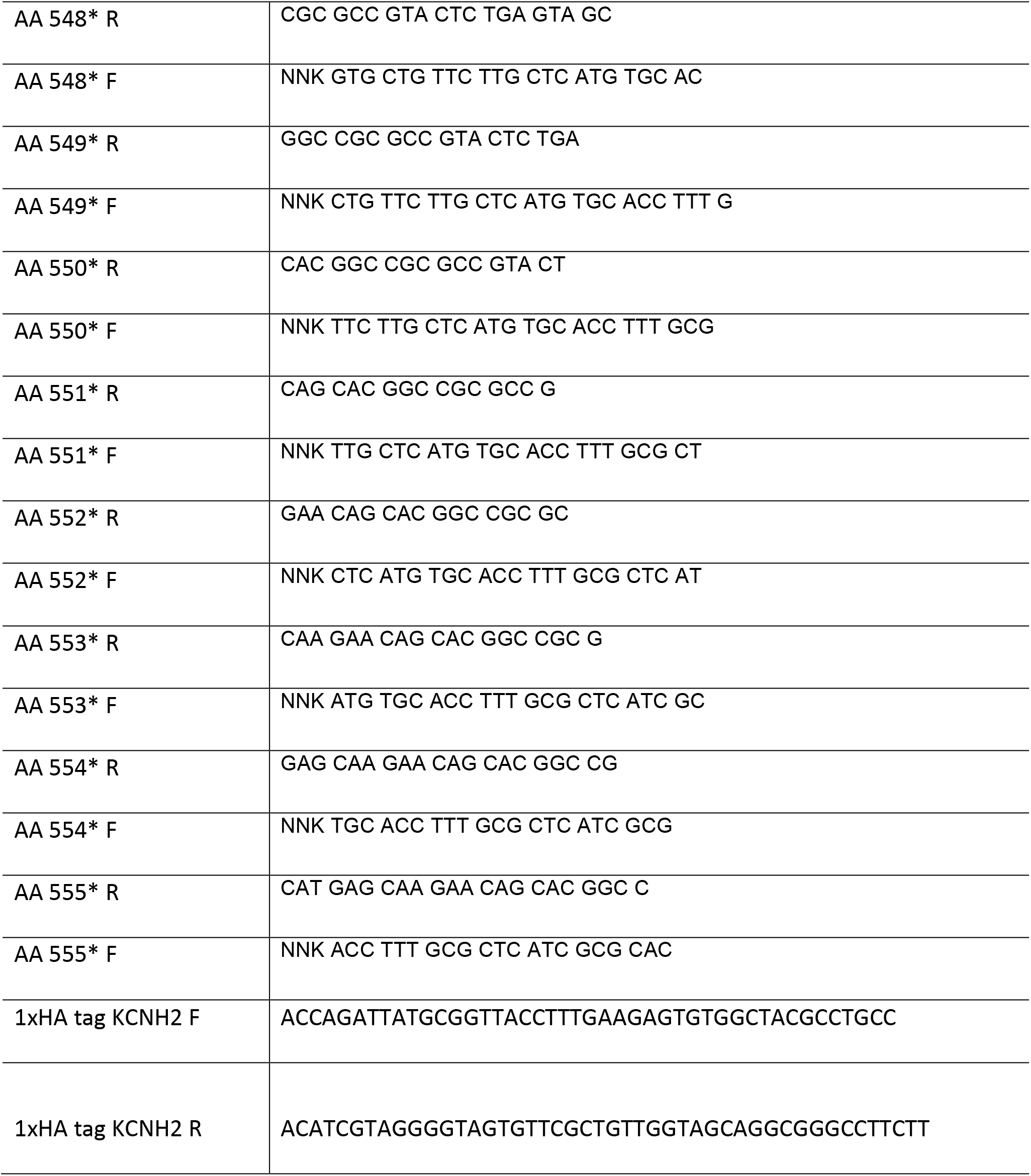
Primers used in this study.

**Table S2:** Trafficking scores for variants observed in individuals to date.

| Variant | DMS score | LQT patients | Unaffected |
| --- | --- | --- | --- |
| A547T | 75 | 0 | 2 |
| V549M | 110 | 0 | 1 |
| F551L | 70 | 0 | 1 |
| L552S | 25 | 90 | 58 |

PROVEAN and PolyPhen-2 predict all variants listed in Table S1 are deleterious; the one exception is F551L which PolyPhen-2 predicts is neutral.

**Figure S1:**
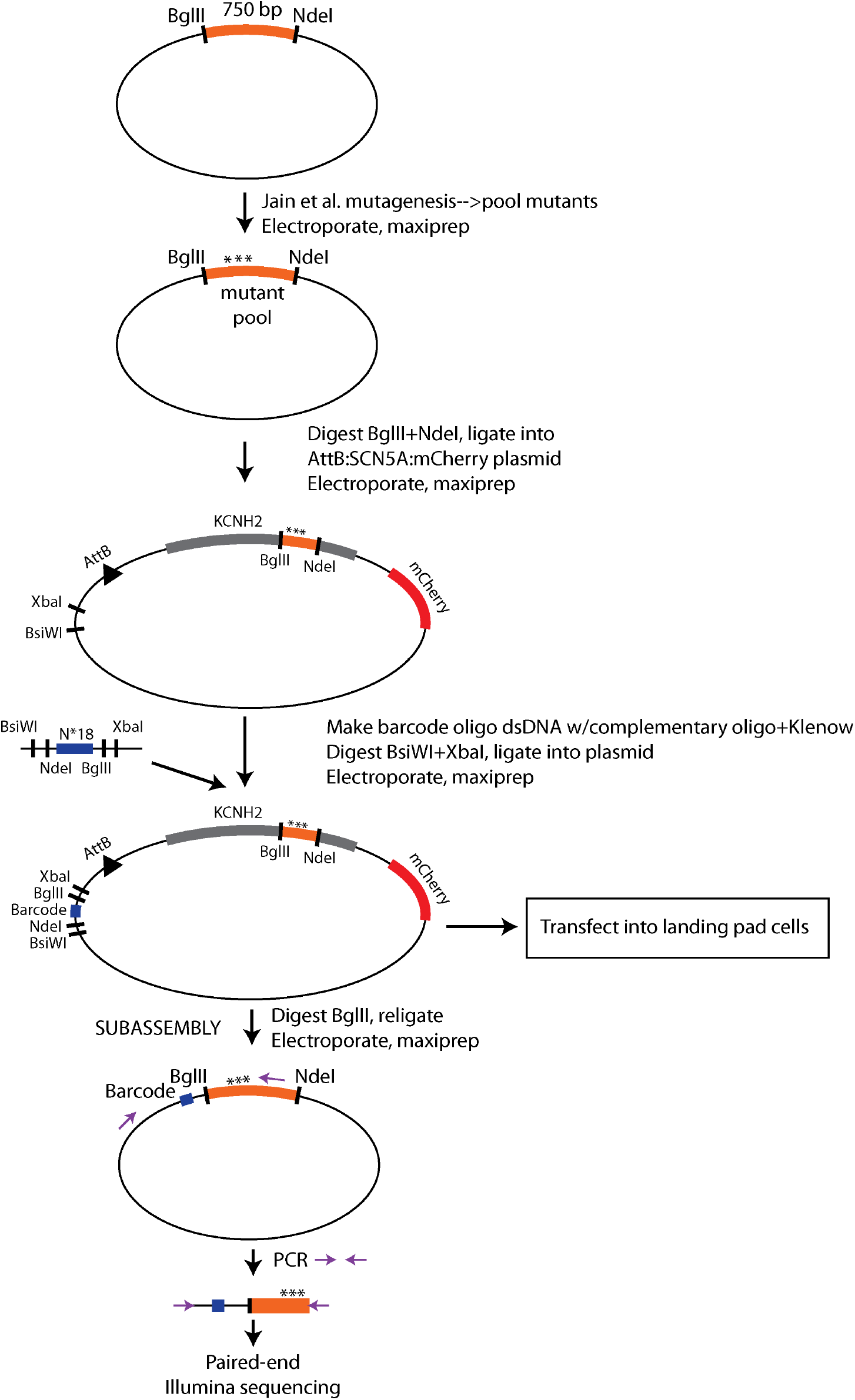
Cloning diagram for Deep Mutational Scan.

**Figure S2:**
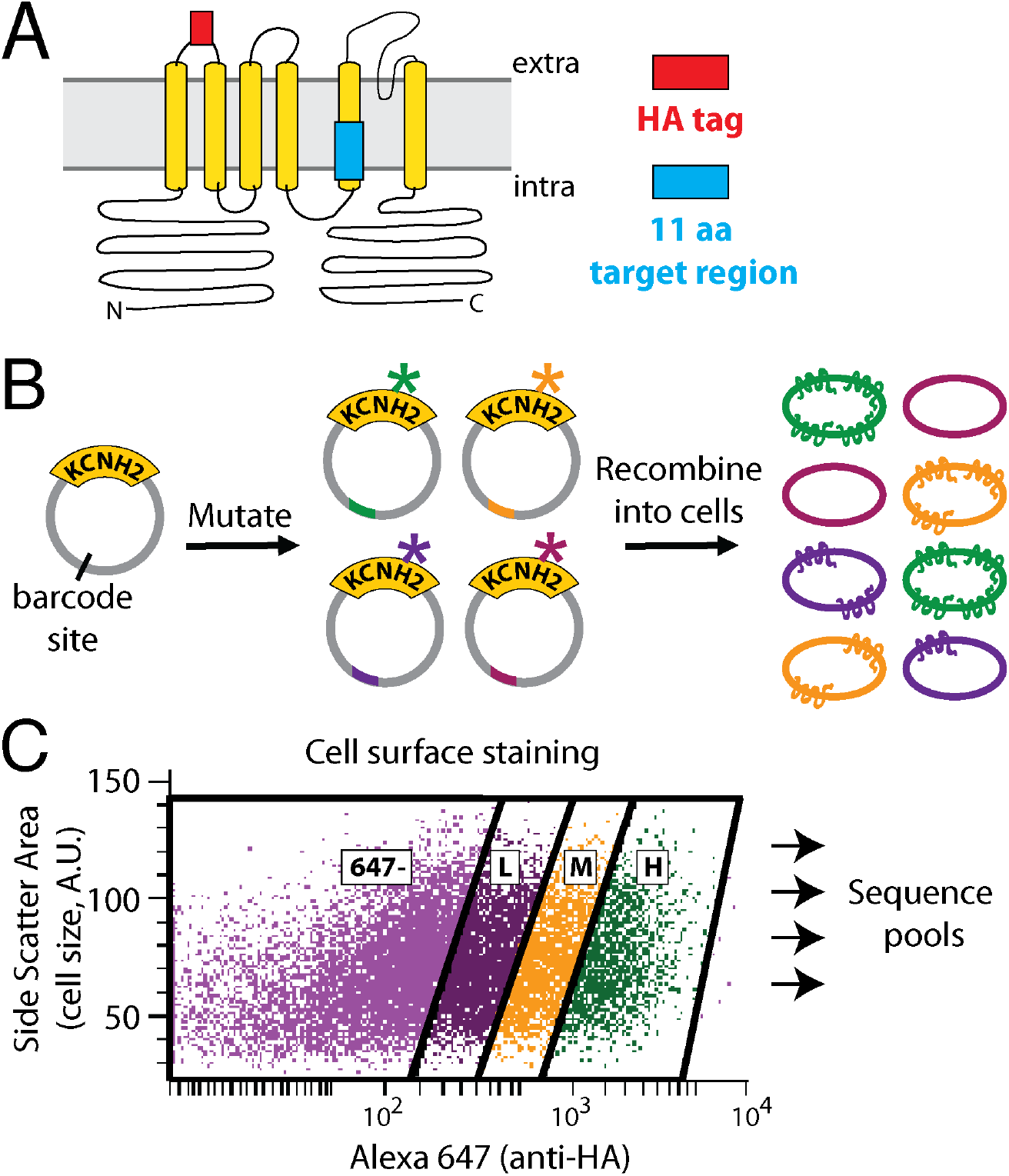
Diagram of Deep Mutational Scan approach. A) Schematic of *KCNH2* showing the location of the extracellular HA tag used for detecting K_V_11.1 localization and the target 11 amino acid region mutagenized in this study (residues 545-555). B-C) Diagram of Deep Mutational Scan method. A mutant *KCNH2* plasmid library was generated and recombined into HEK293 cells. Cells were stained with an Alexa 647-conjugated anti-HA antibody and flow sorted into 4 groups based on their level of Alexa 647 staining. “647-” indicates cells without staining for Alexa 647. Side scatter area is a measure of cell size (arbitrary units). “L”, “M”, and “H” indicate cells with a low, medium, or high level of staining for Alexa 647, respectively. These flow-sorted pools of cells were recovered and Illumina sequenced.

**Figure S3:**
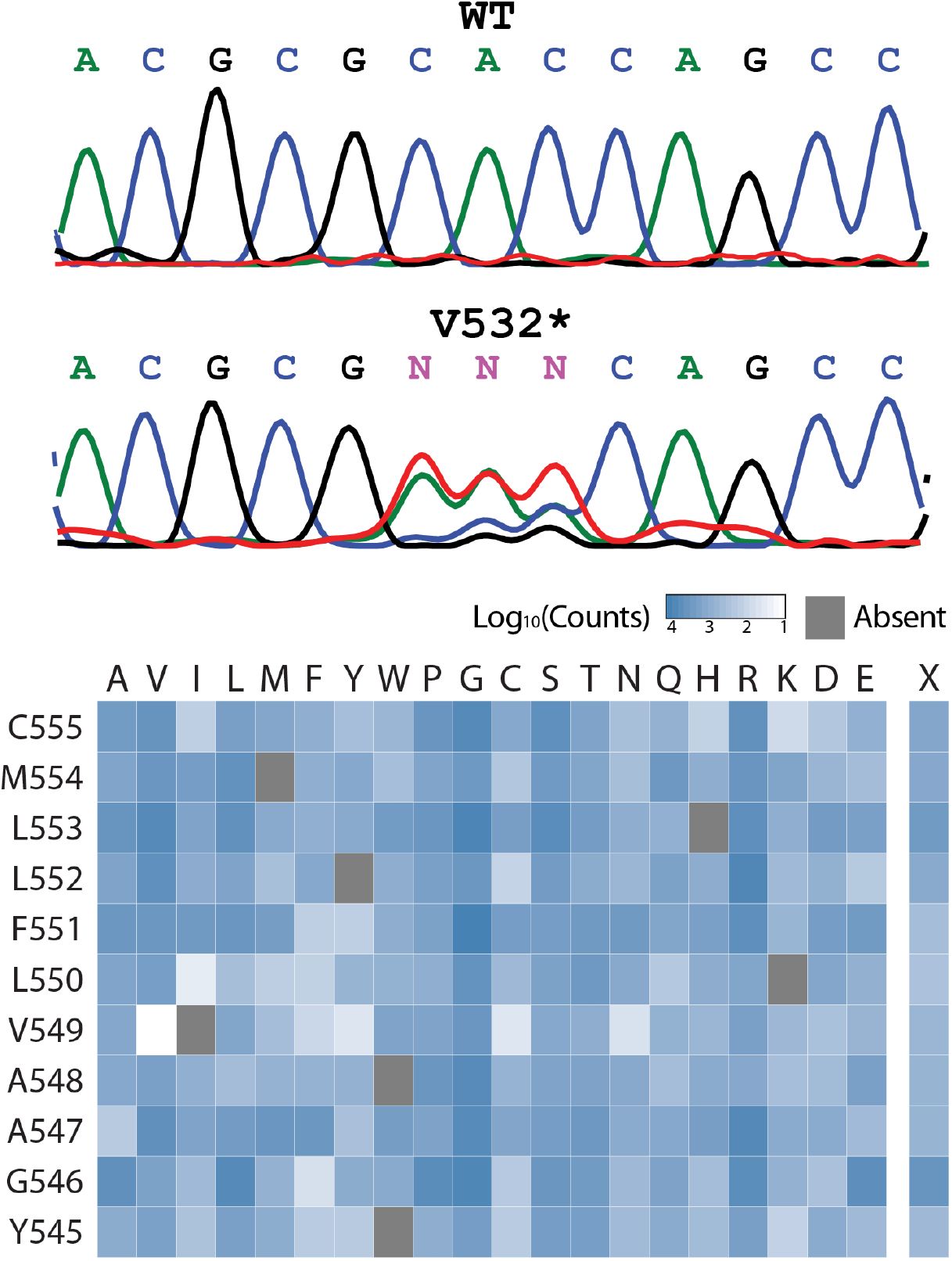
Single codon mutagenesis and abundance of individual amino acid substitutions in mutagenized *KCNH2* library. Top) Mutagenesis of a 1 amino acid region. Sanger sequencing of wildtype and the mutagenized pool indicates a distribution of variants at this position. Bottom) Mutagenesis of the target 11 residue region. Heatmap indicates the abundance of each variant (log_10_ of counts). Grey indicates variant was absent in the library.

**File S1. DMS scores for all variants.**

